# Internally quenched fluorogenic probe provides selective and rapid detection of cathepsin L activity

**DOI:** 10.1101/2020.03.28.012708

**Authors:** Kelton A. Schleyer, Ben Fetrow, Peter Zannes Fatland, Jun Liu, Maya Chaaban, Biwu Ma, Lina Cui

**Affiliations:** Department of Medicinal Chemistry, College of Pharmacy, University of Florida, Gainesville, FL 32610, USA; Department of Chemistry and Chemical Biology, Comprehensive Cancer Center, University of New Mexico, Albuquerque, NM 87131, USA; Department of Chemistry and Biochemistry, Florida State University, Tallahassee, FL 32306, USA

**Author notes:** Correspondence should be addressed to L.C.

## Abstract

Cathepsin L (CTL) is a cysteine protease that demonstrates upregulated activity and/or altered trafficking during disease states such as cancer. The overlapping substrate specificity of cathepsin family members makes selective detection of activity from a single cathepsin difficult, and CTL activity is particularly difficult to parse from its close homologue CTV and the ubiquitous CTB. Despite this, screening campaigns have explored the extended chemical space in the cathepsin binding sites and identified unique substrate structures that offer selectivity for one enzyme over others. In this vein, we present CTLAP, a fluorogenic probe that is rapidly activated by CTL and displays good selectivity over CTB and CTV, the closest competing analytes for CTL activity probes. CTLAP exhibits intrinsically low background fluorescence, which we attribute to possible self-quenching mechanisms. CTLAP demonstrates markedly higher turn-on ratios (24-fold) and moderately improved enzyme selectivity compared to Z-FR-AMC (10-fold turn-on ratio), a commercially available CTL-selective probe commonly used to detect CTL activity in mixed samples. Optimum selectivity for CTL is achieved within 10 min of incubation with the enzyme, suggesting that CTLAP is amenable for rapid detection of CTL, even in the presence of competing cathepsins.

## Introduction

The cathepsin family of lysosomal proteases includes 16 members associated with various pathological conditions^[1]^ including cancer^[2]^. Although their redundancy in certain contexts has been described^[3]^, their distinct tissue distribution^[4–7]^ and association with disease^[1]^ creates a need for specific detection or inhibition^[2]^ of a single cathepsin useful as a biomarker for a specific disease^[8–10]^ or cancer^[11–14]^. For example, CTL is involved in cancer progression by degrading the extracellular matrix (ECM) directly^[15–16]^ while also activating other ECM-degrading enzymes to promote an aggressive phenotype^[7, 17]^. Cathepsin L levels in serum and urine have been shown to increase in cancer patients compared to healthy patients, and tends to correlate with tumor grade, invasive potential, and metastatic spread^[18–23]^. Selective detection of cathepsin L activity is crucial also because it correlates directly with the consequences of the enzymatic activity in question. This information does not always directly correlate with quantified expression at the protein^[24]^ or mRNA level^[25–26]^, including the reduction of CTL activity by the addition of inhibitors^[27]^.

Methods to detect CTL activity typically use the fluorogenic substrate **Z-FR-AMC**(Figure 2) or similar probes which exhibit off-target activation by CTB or other proteases^[28]^. In some cases, removing off-target detection has required inclusion of an exogenous inhibitor^[9, 11, 29]^ or pre-incubation under harsh conditions (4 M urea for 30 min)^[30]^ to deactivate competing enzymes (namely CTB) before detecting activity. These concerns are mitigated with highly selective probes, which have been discovered for CTL activity by screening combinatorial substrate libraries^[31]^, typically requiring generation and testing of thousands to hundreds of thousands of compounds^[32–33]^. Another strategy to develop selective cathepsin L probes has been the ‘reverse-design’ strategy^[34–35]^ which utilizes the information gained from selective inhibitor design and medicinal chemistry optimization to translate a selective inhibitor scaffold into an imaging probe^[36–38]^. Instead of undertaking large-scale screening efforts, investigation of successful inhibitor development can take advantage of previously established SAR relationships^[34]^ and work with already known selective scaffolds to diversify their application. The efficacy of this scaffold repurposing strategy has been demonstrated for targeting cathepsins^[34–35, 39]^ and MMPs^[36–38]^ among other targets. Our work reveals that while reverse-design can retain the selectivity and other properties of the parent inhibitor, it can also result in the discovery of emergent properties that further improve the nature of the imaging probe created. Use of an inhibitor scaffold bearing a benzyl-thiophene group enabled selective detection of CTL while also providing an internal quenching mechanism of the AMC fluorophore in the probe, resulting in a greater turn-on ratio (TOR) compared to commercial standard CTL probe **Z-FR-AMC** (Figure 1).

**Figure 1.**
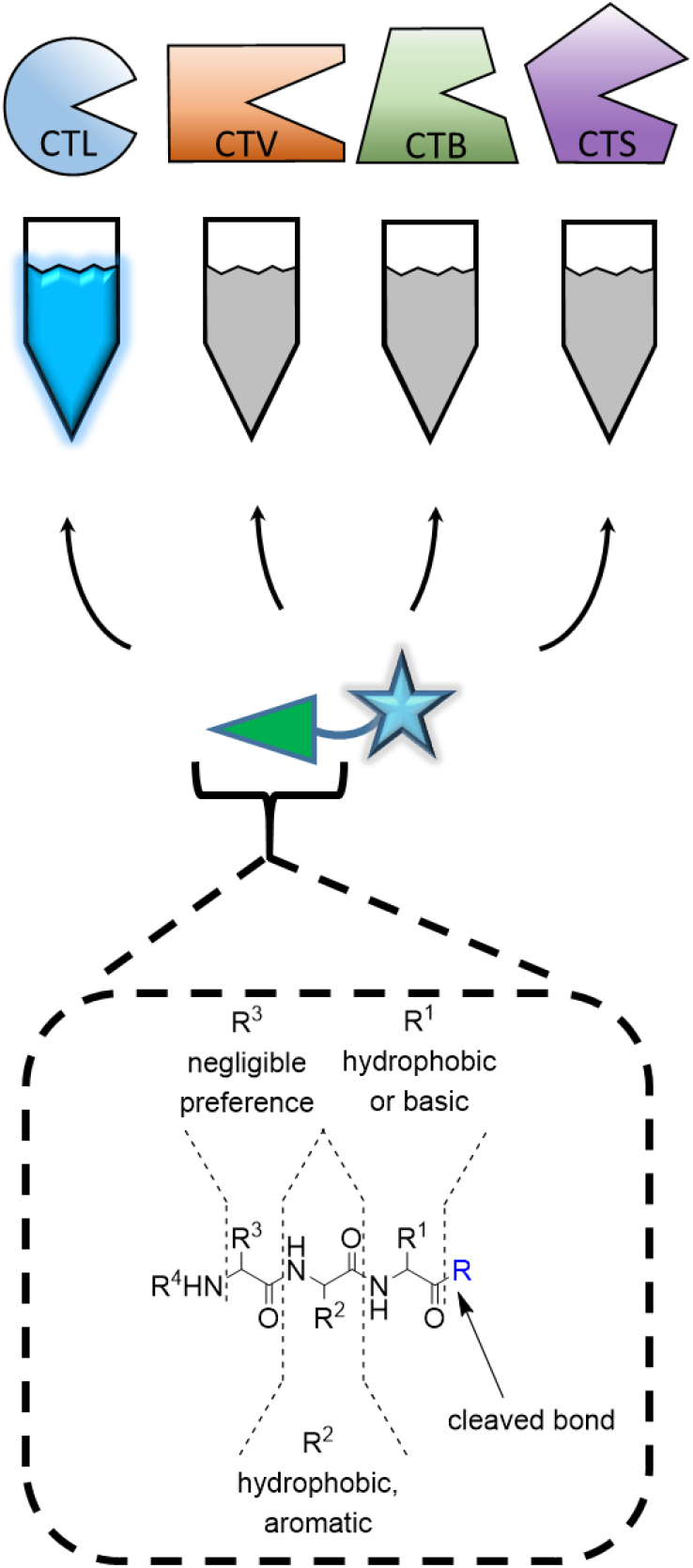
Substrate preference and inhibitor structure (bottom) inform probe design (center), leading to selective detection of the target enzyme, cathepsin L (CTL) over related cathepsins (CTV, CTB, CTS).

**Figure 2.**
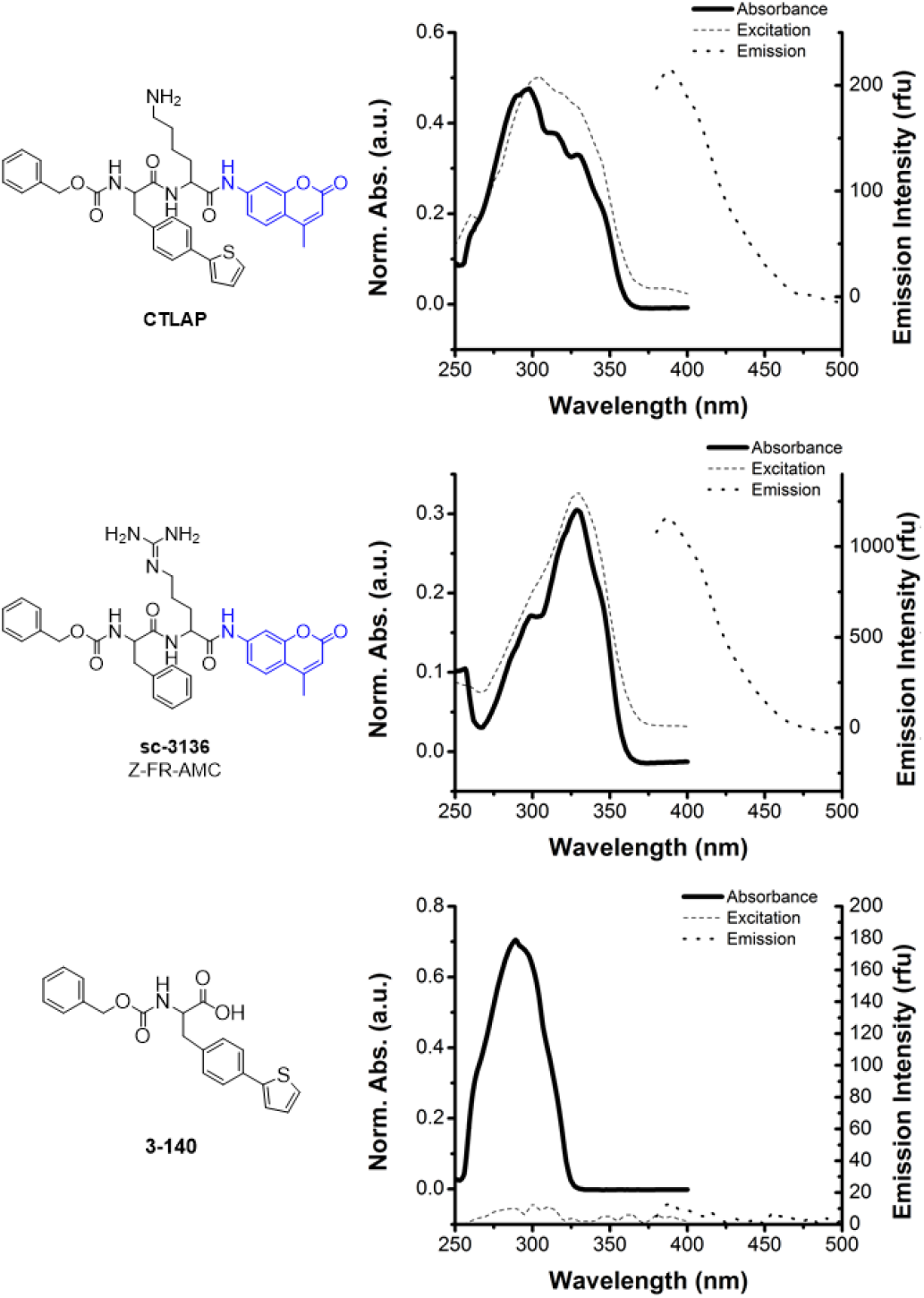
Structures of compounds examined in this work, along with their absorption (solid line), emission, (large dot), and excitation (small dot) spectra. The baseline emission and excitation intensities for **3-140** were included for the sake of completion (see Supporting Discussion).

## Results/Discussion

In seeking attractive inhibitor scaffolds, one notable result in the literature was a benzyl-thiophene residue that provided selectivity for CTL over multiple related cathepsins^[39]^. While examination of the binding pockets elicits no obvious explanation for the selectivity provided by this residue, it appears to be a privileged CTL-selective scaffold even among extended aromatic groups^[39]^. As aromatic residues present in fluorescent probes have been shown to quench the onboard fluorophore^[40]^, this benzyl-thiophene residue held potential to enhance the sensitivity of a fluorogenic probe by reducing background emission, while also retaining the selectivity profile for CTL it showed as an inhibitor. We attached the fluorophore 7-amino-4-methylcoumarin (**AMC**) for direct comparison to **Z-FR-AMC**; a labile linker between the fluorophore and inhibitor scaffold was not included, as such linkers have been shown to shift selectivity away from CTL and toward CTB^[41]^. The resulting cathepsin L-activable probe (**CTLAP**) is shown in Figure 2. The selectivity of **CTLAP** was judged by the turn-on ratio (TOR) of probe emission generated by CTL or other cathepsins (Figure 3, Supporting Figures S1, S2). With **CTLAP**, CTL and CTV are the only enzymes that generate a TOR greater than 2 within 10 min; CTB (the most abundant and ubiquitous cathepsin with activity similar to CTL) only achieved this benchmark at pH 5.0 (ref ^[42]^), indicating that pH 6.5 (ref ^[43]^) is more suitable for CTL-selective detection by **CTLAP**(Supporting Table S1 and Supporting Figure S1). Normalizing CTL signal over time (Figure 3b,c) revealed that in the first 10 min, **CTLAP** generates only 5-10% off-target signal from CTB present at the same concentration as CTL. CTV signal remains below 20% at equal enzyme concentration, but its isolated distribution in the thymus and testes makes its interference less likely than that of the ubiquitous CTB^[5–6]^. These results are only half the interference generated using **Z-FR-AMC**(15-20% background from either CTV or CTB alone), indicating **CTLAP** has greater selectivity for CTL (Supporting Figure S3 and Supporting Table S1). Examining the TOR over the first 1 h incubation also revealed a larger TOR for **CTLAP** than for **Z-FR-AMC** throughout the assay (Supporting Figure S2). In fluorogenic assays, where the readout is fluorescence intensity, a greater TOR means a greater overall SNR for the assay, lending robustness to the assay’s Z-score and viability for HTS applications^[44]^. Generating high TOR/SNR is a major consideration in fluorogenic probes and constitutes a major design strategy^[45]^. The larger TOR of **CTLAP** was attributed to its 6-to 8-fold lower background signal (Figure 4a). With data showing that the absorbance of **CTLAP** is linear within the concentrations used in the assays and the emission of **Z-FR-AMC** is attributed completely to the caged **AMC** moiety (see Supporting Discussion), we hypothesized the lower background intensity of **CTLAP** was due to quenching of the fluorophore by the benzyl-thiophene residue.

**Figure 3.**
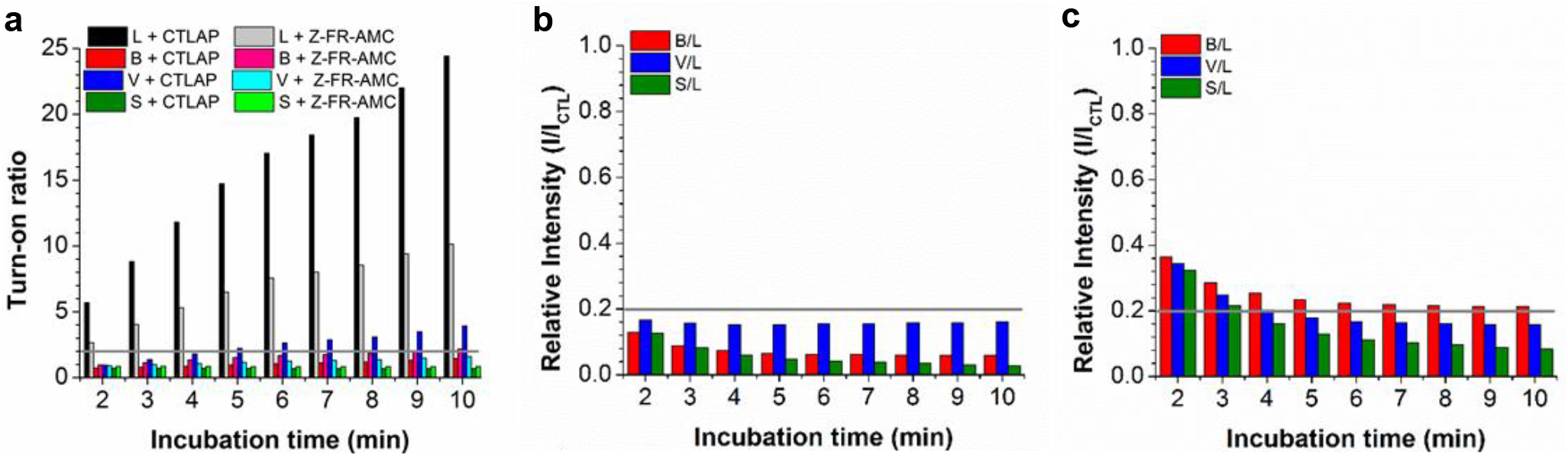
Selectivity of **CTLAP** and **Z-FR-AMC** for cathepsins. (a) Turn-on ratio of probes within 10 min, pH 6.5. Relative signal intensity generated by competing cathepsins with (b) **CTLAP** and (c) **Z-FR-AMC** at pH 6.5. CTL signal was normalized to 1.0 at each time point; horizontal grey line marks 0.2 relative signal.

**Figure 4.**
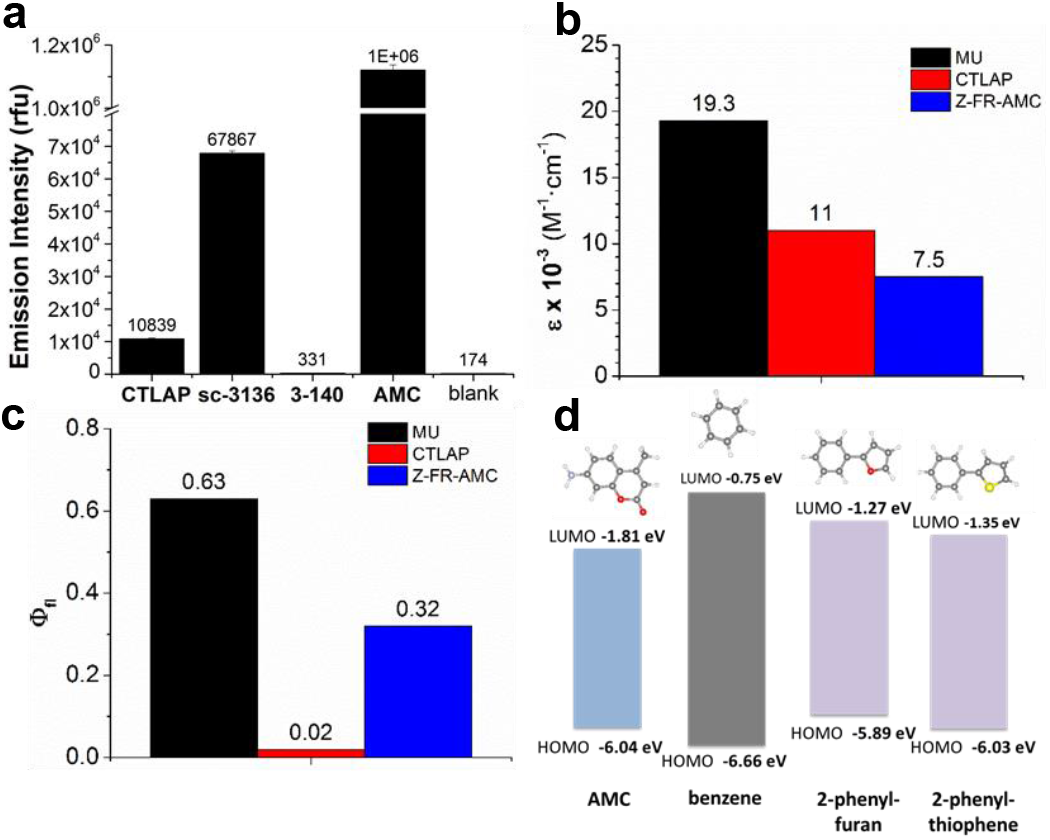
Photophysical data of **CTLAP** and **Z-FR-AMC**. (a) Background emission intensity of 5 μM compound in cathepsin reaction buffer, pH 5.0. Measured (b) molar attenuation coefficient and (c) quantum yield of compounds, standardized against methylumbelliferone (MU). (d) DFT calculations of HOMO and LUMO energies for AMC and aromatic residues of probes.

The measured molar attenuation coefficient of **CTLAP** was greater than that of **Z-FR-AMC**(Figure 4b), however the measured quantum yield of **CTLAP** was significantly lower than that of **Z-FR-AMC**(Figure 4c). As the benzyl-thiophene residue has higher-energy absorption than the **CTLAP** emission wavelength (Figure 2), the possibility of quenching by Forster Resonance Energy Transfer (FRET) from the fluorophore to benzyl-thiophene was considered unlikely. As aromatic amino acid residues have been shown to quench fluorophores^[40]^, we performed DFT calculations to identify the highest occupied molecular orbital (HOMO) and lowest unoccupied molecular orbital (LUMO) of the attached **AMC** fluorophore, as well as the benzyl-thiophene ring of **CTLAP** and the benzyl ring of **Z-FR-AMC**(Figure 4D and Supporting Figure S5). The HOMO-LUMO gap of the extended benzyl-thiophene unit closely matches that of the **AMC** fluorophore, supporting the possibility of energy transfer between the two residues in **CTLAP**. A similar overlap is observed for benzyl-furan, a close analog of the extended aromatic system, suggesting that the two-ring structure is crucial for achieving this similar energy gap. In contrast, the benzyl residue of **Z-FR-AMC** has a notably larger HOMO-LUMO gap, suggesting less potential for energy transfer away from the excited AMC residue, resulting in more frequent emission and higher background signal.

These results support the hypothesis that the unique structure of **CTLAP** reduces the quantum efficiency of the attached **AMC** fluorophore before activation by CTL, an effect not observed in **Z-FR-AMC**.

## Conclusion

Imaging enzyme activity has involved various molecular design strategies to provide selective signal and good contrast^[46]^. Fluorescent probes, including ABPs^[47]^ commonly produce contrast using internal quenching mechanisms, such as bundling reporters together as a polymer probe or attaching a quencher molecule onto the probe structure^[45]^. Upon removal by the target enzyme activity the fluorescent signal is restored, producing high contrast^[46–47]^. Probes lacking such quenching mechanisms provide bright labeling but poor contrast until inactive probes are washed out of the sample^[31]^, compromising their use in rapid or non-invasive procedures such as HTS and clinical imaging. Here we present **CTLAP**, a fluorogenic probe possessing a benzyl-thiophene moiety that contributes to the selectivity of the probe toward CTL while also internally quenching the onboard fluorophore, thereby creating greater contrast and improved detection sensitivity. The increased sensitivity is attributed to a low background emission from the compound, likely due to electron transfer from the excited AMC reporter to the unique benzyl-thiophene residue in the substrate sequence. **CTLAP** exhibits optimal selectivity for CTL within 10 min of incubation time, making it amenable to rapid detection of CTL even in the presence of other cathepsins. Reverse-design has provided a probe scaffold that retains selectivity for CTL while providing greater detection sensitivity by quenching the fluorophore prior to release by the target enzyme. Future work will involve optimizing conditions for applying CTLAP to detection of CTL in clinical samples, cell lysate, and related applications.

## Acknowledgements

This work was supported by research grants to Prof. L. Cui from the University of Florida (UF Startup fund), the National Institute of General Medical Sciences of National Institutes of Health (Maximizing Investigators’ Research Award for Early Stage Investigators, R35GM124963), the Department of Defense Career Development Award (W81XWH-17-1-0529). We are grateful to the support from the University of New Mexico (UNM), the UNM Comprehensive Cancer Center and the National Cancer Institute of the United States (P30CA118100). NMR spectra were collected in part from the NMR Facility, Department of Chemistry and Chemical Biology, UNM, and from the Department of Medicinal Chemistry, College of Pharmacy, University of Florida (UF). Mass spectrometry services were provided in part by the Mass Spectrometry Facility, Department of Chemistry and Chemical Biology, UNM, and from the Mass Spectrometry Research and Education Center, Department of Chemistry, UF (NIH S10 OD021758-01A1).

## Supporting Information Available

The following Supporting Information is available: Text describing general methods and instruments; Figure S1, turn-on ratio of probes at pH 5.0; Figure S2, Turn-on ratio of probes at pH 6.5 and 5.0 for 70 min incubation; Figure S3, relative turn-on ratio of compounds at pH 6.5 and 5.0 for 70 min incubation; Table S1, timepoint of lowest interference from competing cathepsins; Figure S4, measurements of molar attenuation coefficient and quantum yield; Table S2, slope and fits of molar attenuation and quantum yield data; Figure S5, calibration curves of compounds; Supporting Discussion addressing alternative explanations for lower background signal of **CTLAP**; chemical synthesis and characterization data.

