## Supporting Information for "Internally quenched fluorogenic probe provides selective and rapid detection of cathepsin L activity"

#### 1. General reagents and instruments:

Solvents and chemicals were purchased from commercial sources and used directly without further purification. Cathepsin L was purchased from Abcam, and Cathepsins B, V, and S were purchased from Sino Biological. Z-FR-AMC was purchased from Santa Cruz Biotechnology, Inc. Flash chromatography was performed manually with Agela Technologies Flash Silica (40-60 µm, 60 Angstroms). HPLC was performed on a Dionex UltiMate 3000 with a pump and an in-line Diode Array Detector (DAD-3000). A reverse-phase C18 (Teledyne, 5 µm, 10 x 250 mm) column was used for analysis and semi-preparation. $^{1}$H NMR spectra were recorded using either a Bruker Avance III 300, Bruker Avance 500, or a Bruker Avance Neo 600, and chemical shifts were reported in ppm with either TMS or deuterated solvents as internal standards (TMS, 0.00; $CDCl_3$, 7.26; MeOD, 3.31). $^{13}$C NMR spectra were recorded at 75.4 or 125.7 MHz, and chemical shifts were reported in ppm with deuterated solvents as internal standards ($CDCl_3$, 77.0; MeOD, 49.15). Absorption, emission, and excitation data were collected using a BioTek Synergy H1 plate reader. UV-vis spectra were collected on a Shimadzu UV-2700 spectrophotometer.

#### 2. Methods

**Enzyme storage and preparation:**

Enzymes were reconstituted in accordance with manufacturer's instructions, then diluted to 2.5 µM in a storage buffer according to a literature method[1]: 100 mM NaOAc/AcOH buffer (pH 5.5), with 2.5 mM DTT and 2.5 mM EDTA, containing 5% v/v glycerol. Cathepsin L stock was prepared as described by a different procedure[2] (100 mM NaOAc/MaOH, 100 mM NaCl, 10 mM DTT, 1 mM EDTA, pH 5.5), to a final concentration of 2.5 µM. The stocks were divided into aliquots and stored at -80 C.

**Cathepsin activity assays:**

All enzyme reactions and assays took place in 100 mM NaOAc/HOAc, 2.5 mM DTT, 2.5 mM EDTA, at either pH 5.0 or 6.5, with final concentration of 5 nM cathepsin and 5 mM **CTLAP** and **Z-FR-AMC**. Fluorescence intensity measured using a BioTek Synergy H1 plate reader, at 350 nm excitation and 445 nm emission unless otherwise noted. For longer experiments (2 to 48 h), plates were incubated at 37 C for the course of the experiment. All data points were prepared in triplicate, and values are reported as the average of three samples; error bars depict standard deviation of the mean, when shown.

**UV-vis and fluorescence spectra of CTLAP and Z-FR-AMC:**

Excitation and emission spectra were collected on solutions of 5 µM **CTLAP** and **Z-FR-AMC** in cathepsin reaction buffer (pH 5.0 or 6.5) using a BioTek Synergy H1 plate reader. Absorbance spectra of compounds (80-200 µM in DMSO) were collected on a Shimadzu UV-2700 spectrophotometer.

**Measurement of molar absorption coefficient and quantum yield:**

To determine quantum yields, standard curves of **CTLAP** and **Z-FR-AMC** were constructed between 1 – 10 µM in carbonate buffer (pH 10). Using a BioTek Synergy H1 plate reader, absorbance at 350 nm was measured, and the fluorescence curve at 350 nm excitation and 380-600 nm emission was measured. The absorbance at 350 nm was graphed against the area under the curve (AUC) of the fluorescence spectrum from 380-600 nm, and the slope of each line was used to calculate the quantum yield according to the equation (1) (see ref [3]):

$$\phi_x = \phi_{st}\left(\frac{Grad_x}{Grad_{st}}\right)\left(\frac{\eta_x^2}{\eta_{st}^2}\right) \qquad (1)$$

Where *st* is a standard reference compound, *x* is an unknown compound to be measured, Φ is the quantum yield, $Grad_x$ is the slope of the plot of integrated fluorescence intensity as a function of absorbance, and η is the refractive index of the solvent. Values for the molar attenuation coefficient and quantum yield were standardized using measured absorption and emission intensity of 4-methylumbelliferone (MU) and its literature quantum yield and molar attenuation coefficient[4].

**Calibration curves of absorbance and emission:**

For AMC emission calibration, solutions of AMC between 0.5-7.5 µM were prepared in cathepsin buffer (pH 5.0, 5.5, and 6.5). Emission calibration curves of **CTLAP** and **Z-FR-AMC** were constructed between 2-16 µM in cathepsin reaction buffer (pH 5.0). For pH-dependence of AMC emission, 5 µM AMC prepared in buffers ranging from pH 3.0 to 11.0. Citric acid (0.1 M) and disodium hydrogen phosphate (0.2 M) buffer was used for pH 3.0, 4.0, 5.0, and 6.0; disodium hydrogen phosphate (0.2 M) and sodium dihydrogen phosphate (0.2 M) was used for pH 7.0 and 8.0; sodium carbonate (0.1 M) and sodium bicarbonate (0.1 M) was used for pH 9.0, 10.0, and 11.0. All measurements were performed using a BioTek Synergy H1 plate reader. All data points were prepared in triplicate, and values were reported as the average of three samples; error bars depict standard deviation of the mean, when shown.

#### 3. Supporting Figures and Tables

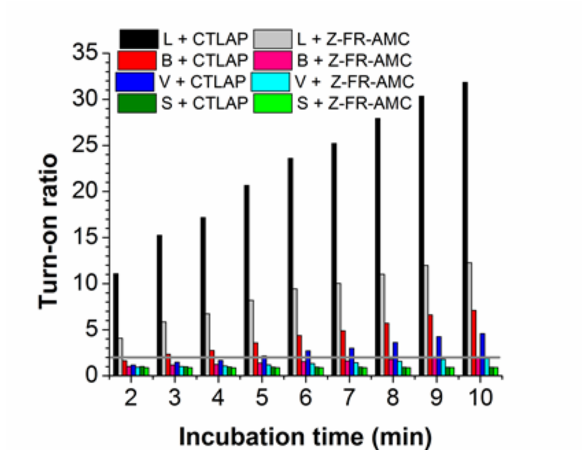

**Supporting Figure S1.** Turn-on ratio (TOR) of **CTLAP** and **Z-FR-AMC** within the first 10 min incubation with CTL, CTB, CTV, and CTS, at pH 5.0. The horizontal grey line marks a TOR of 2.

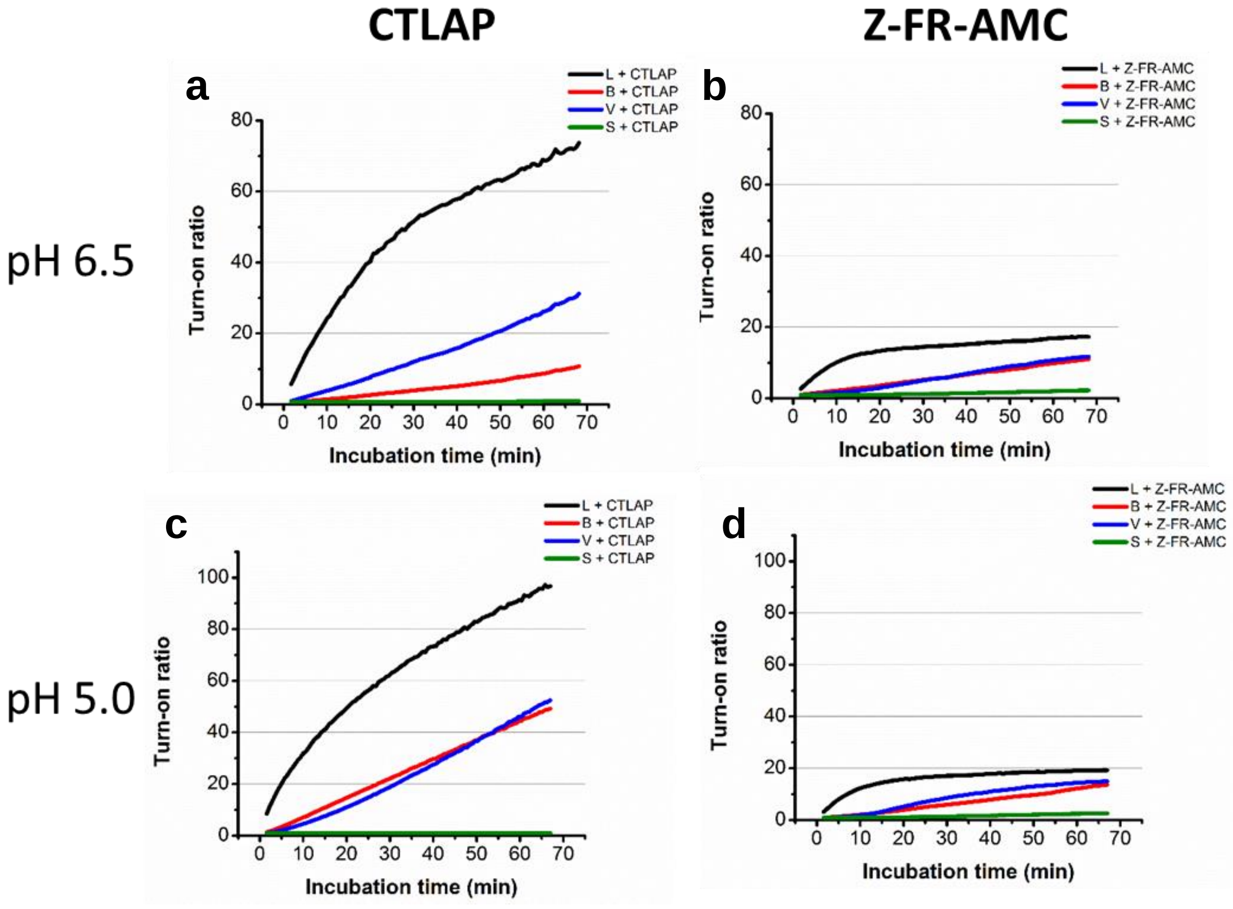

**Supporting Figure S2.** Turn-on ratio of (a, c) **CTLAP** and (b, d) **Z-FR-AMC** incubated with cathepsins over 70 min, at (a, b) pH 6.5 or (c, d) pH 5.0.

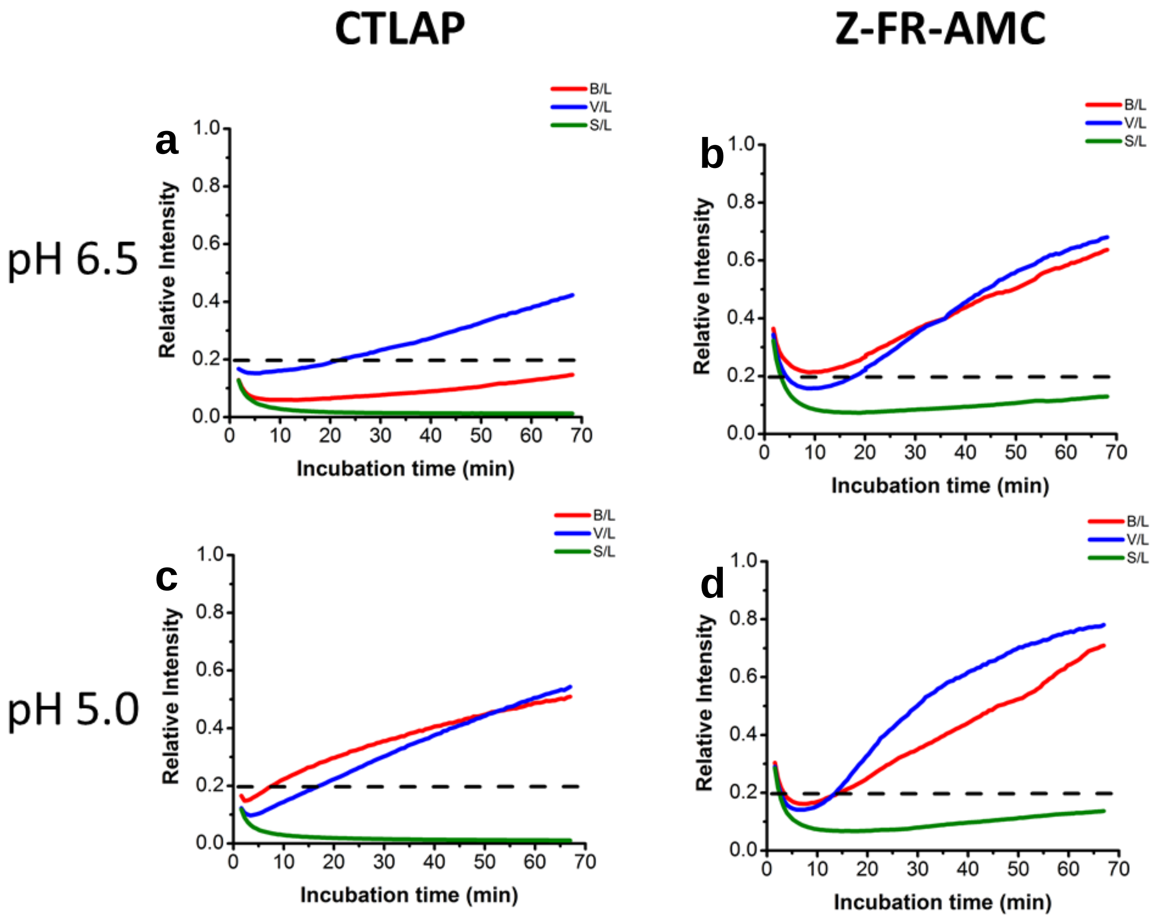

**Supporting Figure S3**. Relative TOR for (a, c) **CTLAP** or (b, d) **Z-FR-AMC** incubated with CTB, CTV, CTS for 70 min. CTL signal was normalized to 1.0 at each time point. The dashed line marks 0.2 relative signal. Incubation with at (a, b) pH 6.5 or (c, d) pH 5.0.

| | CTLAP | | sc-3136 | |
|---|---|---|---|---|
| pH | 5.0 | 6.5 | 5.0 | 6.5 |
| CTB | 15% | 6% | 16% | 21% |
| CTV | 10% | 15% | 14% | 16% |
| CTS | 1% | 1% | 7% | 7% |
| B+V+S | 31% | 25% | 39% | 46% |
| time (min) | 3.4-4.0 | 10.2-11.4 | 7.6 | 11.4 |

**Supporting Table S1.** Lowest relative TOR measured for each competing enzyme within the first 10 min incubation, extracted from Supporting Figure S3. The timepoint giving the lowest interference is also provided in the bottom row.

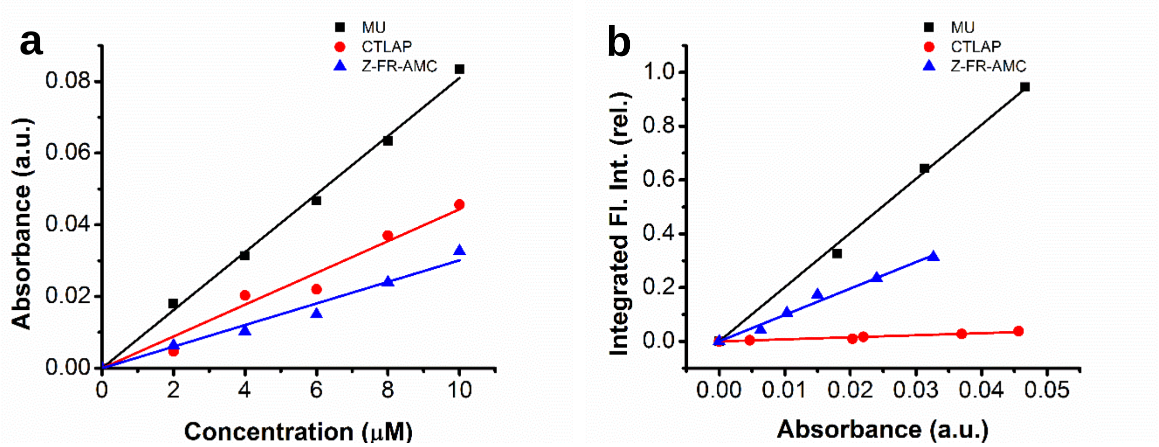

**Supporting Figure S4.** Measurements of (a) molar attenuation coefficient and (b) quantum yield of **CTLAP** and **Z-FR-AMC**, standardized against methylumbelliferone (MU). The slopes and fits of the curves are shown in Supporting Table S2.

| Compound | ε | | $\phi_F$ | |
|---|---|---|---|---|
| | slope | $R^2$ | slope | $R^2$ |
| MU | 0.008 | 0.998 | 19.318 | 0.997 |
| CTLAP | 0.004 | 0.986 | 0.761 | 0.978 |
| sc-3136 | 0.003 | 0.988 | 9.827 | 0.993 |

**Supporting Table S2**. Slopes and fits of molar attenuation coefficient and quantum yield measurements of compounds, from Supporting Figure S4.

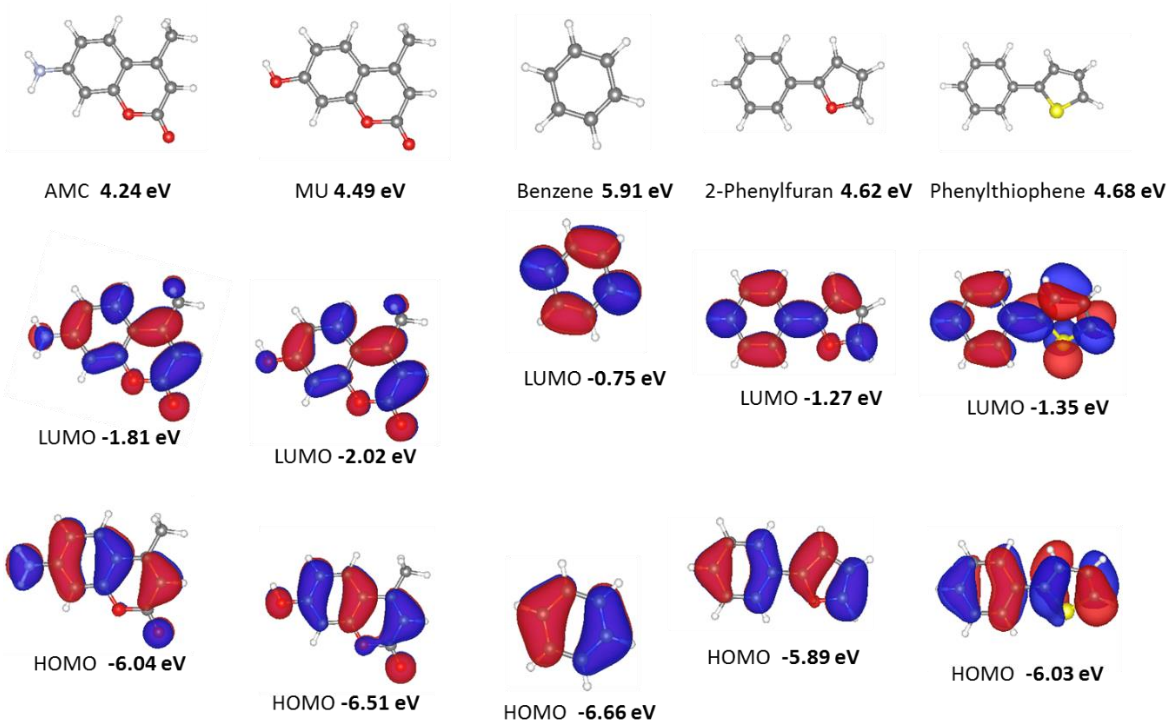

**Supporting Figure S5.** HOMO and LUMO electron densities calculated for 7-amino-4-methylcoumarin (AMC), methylumbelliferone (MU), benzene (representing **Z-FR-AMC**), 2-phenylfuran, and 2-phenylthiophene (representing **CTLAP**).

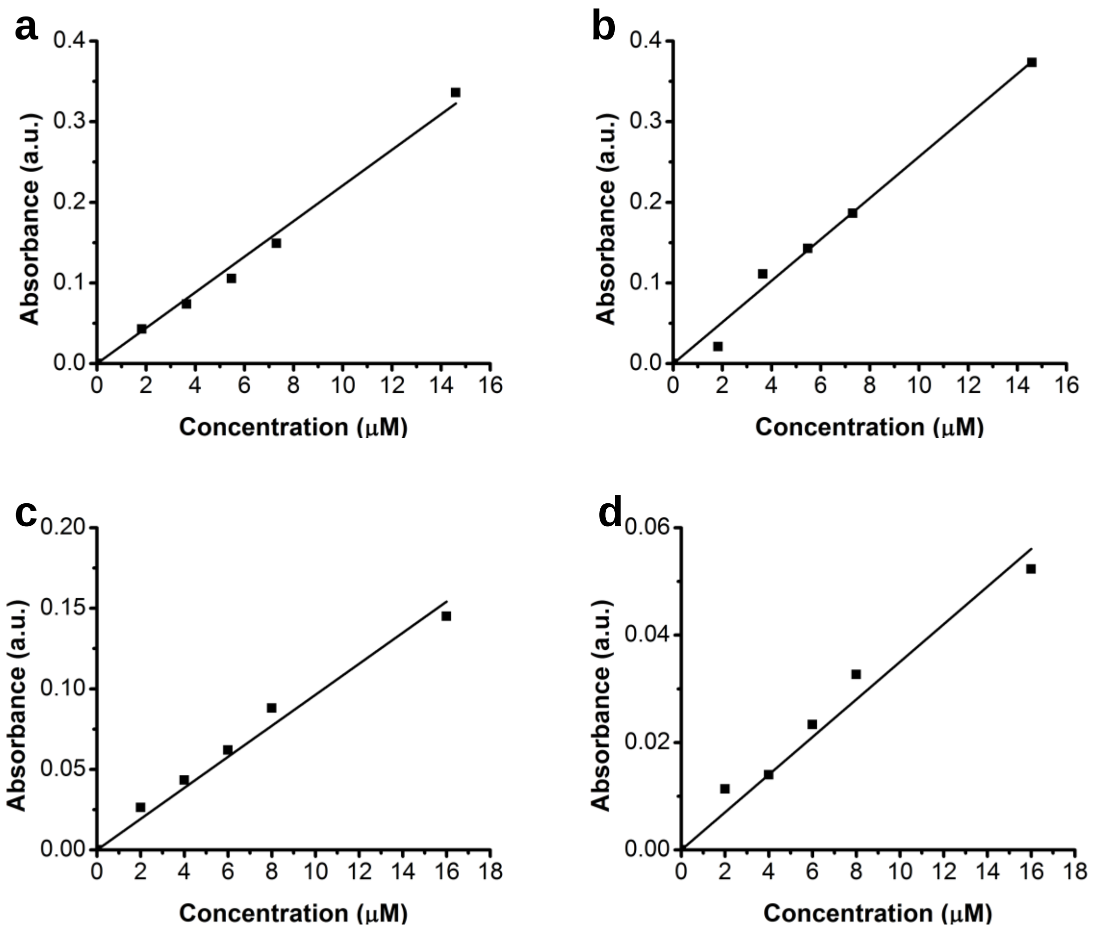

**Supporting Figure S6**. Calibration curves of (a, b) **CTLAP** and (c, d) **Z-FR-AMC** absorbance between 2 μM and 16 μM in cathepsin reaction buffer, pH 5.0. Absorbance measured at (a, c) 330 nm and (b, d) 350 nm.

### 4. Supporting Discussion

**CTLAP absorbance has linear relationship with concentration near assay concentration.** In our assays the working concentration of compounds was calculated using a calibration curve of the compound absorbance at 330 nm, across the concentration range tested. All enzyme assays used 350 nm excitation, 445 nm emission wavelengths. The observed lower background intensity for **CTLAP** could be explained by three general hypotheses: (1) the absorbance values used in calibrating the compound stock concentration were inaccurate; (2) the measured emission intensities of the compounds were inaccurate, reporting a consistently higher value for **CTLAP** in spite of the compound being at equal concentration to that of **Z-FR-AMC**; (3) the quantum yield of **CTLAP** is lower than that of **Z-FR-AMC**, resulting in lower emission intensity at equal concentration. To test hypothesis (1), that the absorbance values of **CTLAP** are inaccurate and result in a lower true concentration of probe used in these assays, we prepared calibration curves of **CTLAP** and **Z-FR-AMC** from 0 – 16 µM in buffer: a good linear increase in absorbance was observed for both **Z-FR-AMC** ($R^2$ = .9781) and **CTLAP** ($R^2$ = .9896) (Supporting Figure S6), suggesting that no significant scattering effects were present in either compound at the concentration used for the enzyme assays (5 µM). Upon incubation with CTL, both compounds are hydrolyzed to generate the free **AMC** reporter; we also measured absorbance of **AMC** from 0-8 µM, and a good fit ($R^2$ = 0.996) was found within this concentration range (data not shown).

We next examined whether another portion of the **CTLAP** structure absorbs 330 nm light, resulting in an artificially high concentration calculation and a true assay concentration lower than that of **Z-FR-AMC**, resulting in reduced emission intensity. We measured UV-vis absorbance spectra of **CTLAP**, **Z-FR-AMC**, **3-140**, and **AMC** between 250 nm and 400 nm and compared the spectral characteristics of the compounds (Figure 2, main text). To establish a standard reference point for the absorption curve of caged **AMC**, we examined the absorption spectrum of **Z-FR-AMC**. The compound exhibits a broad absorbance band with a maximum at 330 nm and a low-energy shoulder peaking around 345 nm. Similarly, **CTLAP** absorbance shows a local maximum at 330 nm with a similar shoulder at 345 nm. However, another band is present at higher energy, bearing a maximum at 300 nm and a low-energy shoulder at 315 nm which slopes down into the **AMC**-regime maximum of 330 nm, still a prominent maximum. As the only prominent structural difference between **CTLAP** and **Z-FR-AMC** is the extended benzyl-thiophene group of **CTLAP**, we examined the absorption spectrum of that moiety alone using compound **3-140** to determine if these new absorption bands in **CTLAP** were from this moiety. The absorption spectrum of **3-140** shows a single broad curve with a maximum at 290 nm and two near-symmetric, gentle shoulders protruding at 265 nm and 315 nm. Visual comparison with **CTLAP** absorption supports the conclusion that the higher energy absorption bands in **CTLAP** come from the benzyl-thiophene structure; thus, the overall absorption of **CTLAP** is the sum of absorption from caged **AMC** and the benzyl-thiophene unit. Significantly, the benzyl-thiophene group alone (compound **3-140**) shows almost negligible absorption at 330 nm (note the excitation spectrum in Figure 2, main text), indicating that this unique structural characteristic of **CTLAP** should not contribute to the absorption measurements taken at this wavelength to calibrate the compound concentration, especially not to an extent causing a 6-fold error in calculated concentration. From this, the absorbance value of **CTLAP** at 330 nm can be solely attributed to the **AMC** portion of the compound, meaning the true and calculated concentration of **CTLAP** in all assay is equivalent to that of **Z-FR-AMC**.

**Emission intensity of CTLAP and Z-FR-AMC is attributed exclusively to the AMC fluorophore**. We next tested hypothesis (2), that the emission intensity measured for **CTLAP** or **Z-FR-AMC** was somehow compromised by the experimental conditions or by the structural differences of the compounds. The presence of free **AMC** impurities in stock solutions of **Z-FR-AMC** would increase the emission measured at 445 nm, causing the greater emission intensity observed for the compound. HPLC traces of probe conversion by CTL (data not shown) indicate no **AMC** peak was detected in the assay wells containing **CTLAP** alone or **Z-FR-AMC** alone; regardless, the presence of free **AMC** would increase the emission intensity when exciting at 350 nm, more than at 330 nm (as 350 nm is closer to the absorption maximum of **AMC**), which is the opposite from what is observed with **Z-FR-AMC**. While emission intensity could be influenced by exogenous emitters present in the solution, this effect could also be produced by the presence of other emitters within the chemical structure of **Z-FR-AMC**. Judging from the spectra, **Z-FR-AMC** emits exclusively from the **AMC** residue and no other portions of the molecule are augmenting the emission intensity. In conclusion, the emission and excitation data indicate that emission from **Z-FR-AMC** can be attributed solely to the caged **AMC** moiety, and no other emitters are present that could be contributing to the greater background emission intensity observed for the compound.

### 5. Chemical Synthesis and Characterization

##### Chemical synthesis:

Compounds used in the study were synthesized by the route outlined in **Scheme S1**.

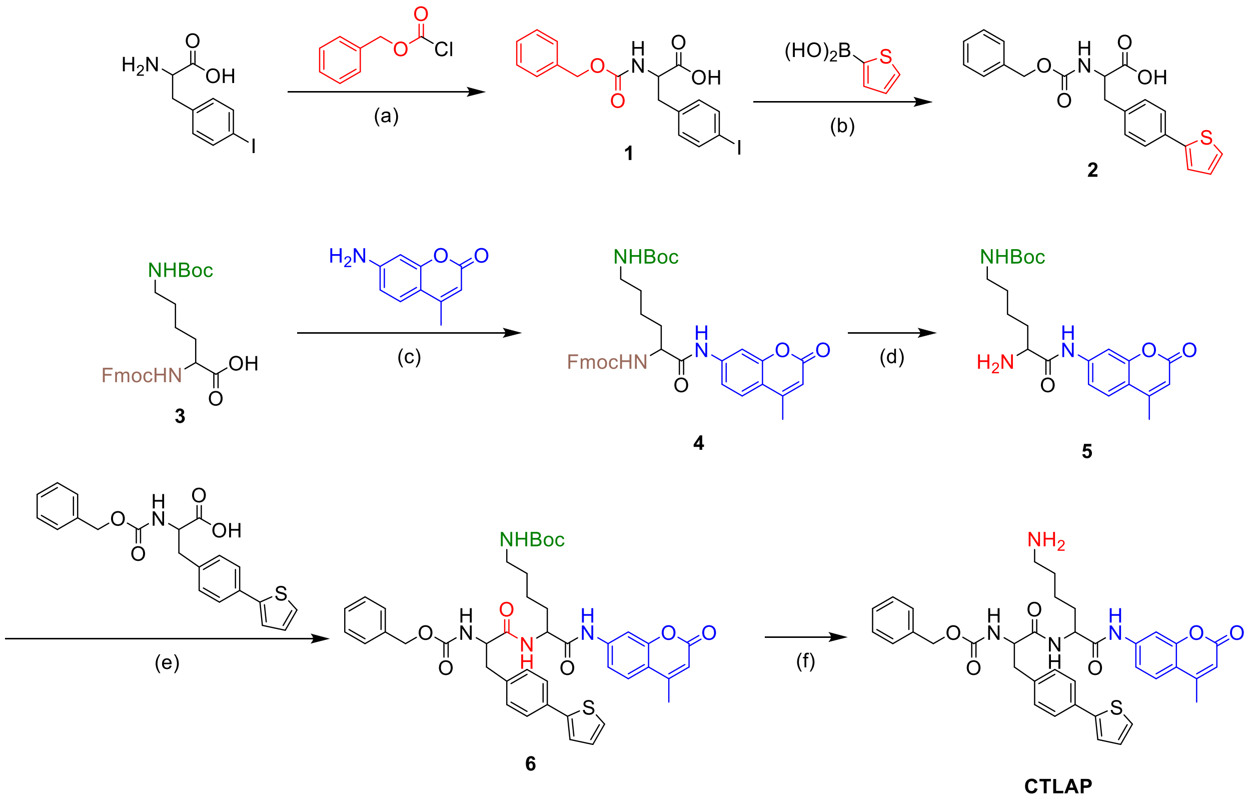

**Scheme S1.** Synthetic scheme of target compound, **CTLAP**. *Reagents and conditions:* (a) benzyl chloroformate, $K_2CO_3$, toluene, $H_2O$, 0ºC -> r.t., 22 h, **82%**; (b) thiophene-2-boronic acid, $Pd(PPh_3)_4$, $K_2CO_3$, $MeCN/H_2O$ (3:1), 80ºC, OVN, **56%**; (c) 7-amino-4-methylcoumarin (**AMC**), $POCl_3$, pyridine, THF (anhydrous), 0ºC, 1.5 h; (d) 5% piperidine/DMF, r.t., 5 min, **65%**; (e) **Z-Phe(thiophene)-OH**, HBTU, HOBt, DIPEA, DMF (anhydrous), r.t., 30 min; (f) 20% TFA/DCM, triisopropylsilane, r.t., 30 min, **quantitative (HPLC)**.

**Characterization of molecules used in the study:**

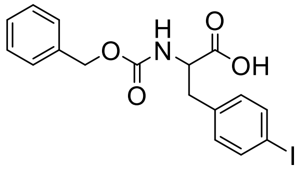

**2-(((benzyloxy)carbonyl)amino)-3-(4-iodophenyl)propanoic acid (1):** $K_2CO_3$ (712 mg, 5.15 mmol, 3.0 eq.) was dissolved in $H_2O$. To the clear solution was added L-phenylalanine (500 mg, 1.72 mmol, 1.0 eq.), and the suspension was stirred and chilled in an ice bath. In a separate container, benzyl chloroformate (366 uL, 2.57 mmol, 1.5 eq.) was diluted with toluene (500 uL, to make 50% solution with benzyl chloroformate) and the solution was added dropwise to the reaction solution. Reaction allowed to gradually return to room temperature overnight, after which TLC indicated one spot ($R_f$ = 0.37; hexanes/ethyl acetate [1:1] with 1.0% acetic acid) that was UV-active (254 nm) and stained negative (faint brown color) with ninhydrin. The reaction solution was diluted with ethyl acetate, acidified with HCl until pH = 1-2, and mixed until any resulting white precipitate dissolved. Extracted with ethyl acetate and residual solids dissolved. The murky organic layer was washed with acidified (HCl) brine, providing a clear organic layer which was subsequently dried over $Na_2SO_4$. Purified by flash column chromatography using isocratic hexanes/ethyl acetate [1:1] with 1.0% acetic acid. Product fractions co-evaporated with toluene to remove trace acetic acid. **82%** yield. $[M+Na]^+$ = 448.0047 m/Z observed; $[M-H]^-$ = 424.0096 observed.

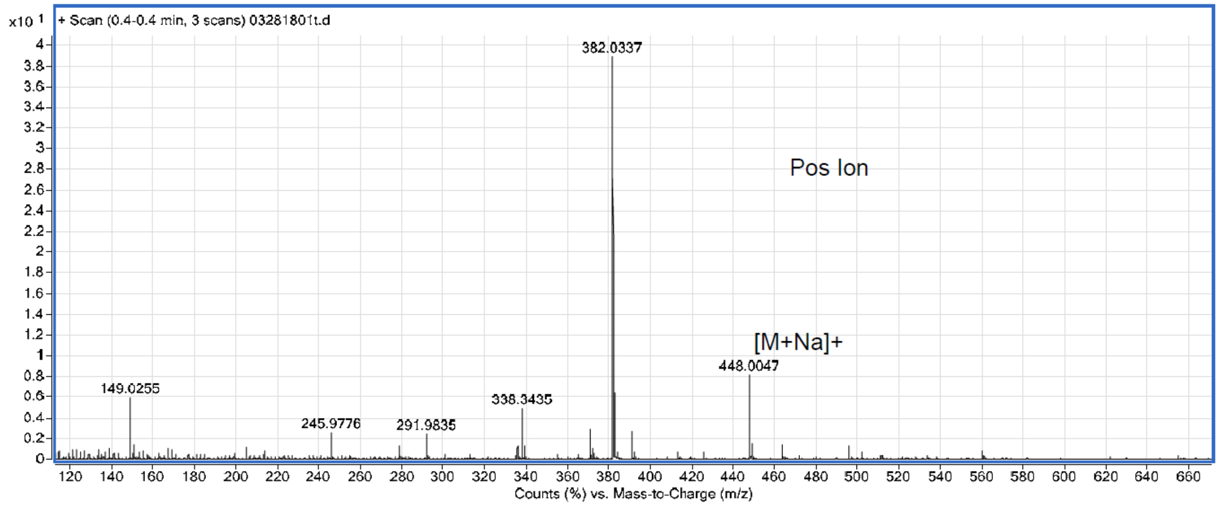

**Supporting Figure S7.** ESI-MS (positive mode) of compound **1**.

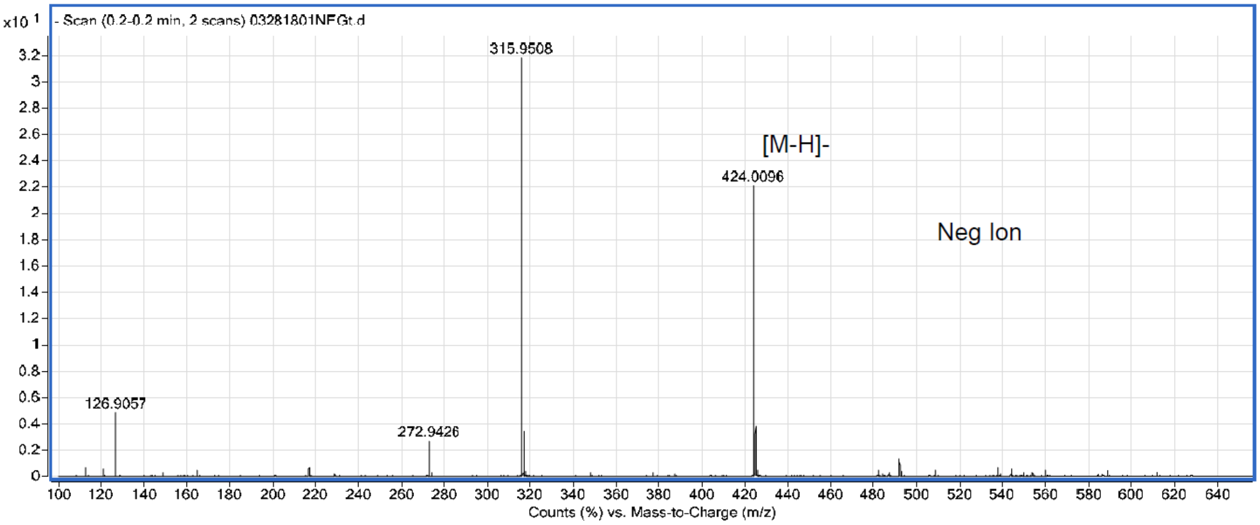

**Supporting Figure S8.** ESI-MS (negative mode) of compound **1**.

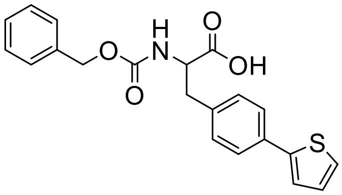

**2-(((benzyloxy)carbonyl)amino)-3-(4-(thiophen-2-yl)phenyl)propanoic acid (2):** Compound **1** (464 mg, 1.09 mmol, 1.0 eq.) and thiophene-2-boronic acid (209 mg, 1.635 mmol, 1.5 eq.) were suspended in MeCN/$H_2O$ (3:1 v/v, 12 mL total). The mixture was stirred and $Pd(PPh_3)_4$ was added to give pale-yellow, fine suspension. $K_2CO_3$ added, reaction flask covered in tin foil, and stirred at 80 C overnight. TLC showed a new, slightly less-polar spot ($R_f$ = 0.59; hexanes/ethyl acetate [1:2] with 1.0% acetic acid) that was UV-active (254 nm) and showed dark blue fluorescence under 302 nm light; also detected by $KMnO_4$ stain. Upon completion, reaction was diluted with ethyl acetate, water, and a small aliquot of 1 M HCl (to insure complete dissolution of product, pH < 3). The bi-layer solution was filtered through Celite to remove palladium, and the funnel was rinsed with ethyl acetate then water to ensure product was in filtrate. The filtrate was transferred to a separatory funnel and extracted with ethyl acetate, then washed with acidified brine. The translucent brown organic layer was dried over $Na_2SO_4$, then concentrated *in vacuo* to afford a viscous brown residue. Purified by flash column chromatography using isocratic hexanes/ethyl acetate (2:1) with 1.0% acetic acid. Brown solid product, **56%**. MS: $[M+H]^+$ calc'd. 382.1105, obs. 382.1067; $[M+Na]^+$ calc'd. 404.0932, obs. 404.1365.

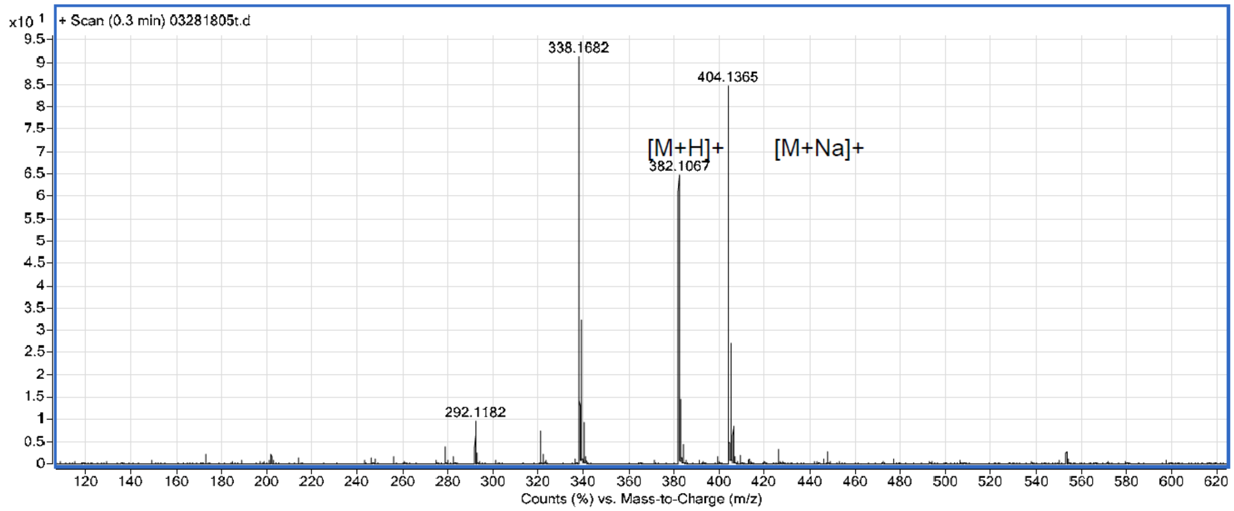

**Supporting Figure S9.** ESI-MS (negative mode) of compound **1**.

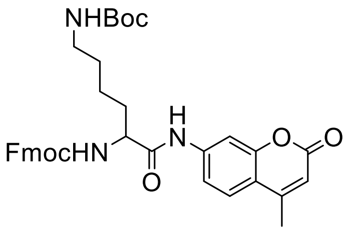

**(9H-fluoren-9-yl)methyl tert-butyl (6-((4-methyl-2-oxo-2H-chromen-7-yl)amino)-6-oxohexane-1,5-diyl)dicarbamate (3):** Fmoc-Lys(Boc)-OH (4.8 mg, 0.01 mmol, 1.0 eq.) and 7-amino-4-methylcoumarin (1.8 mg, 0.01 mmol, 1.0 eq.) were added to an $N_2$-charged flask. THF (1 mL) added to give clear, homogeneous solution, which was stirred and chilled in an ice bath. Pyridine (8.3 uL, 0.10 mmol, 10.0 eq.) added to give pH = 6. $POCl_3$ (4 uL, 0.043 mmol, 4.2 eq.) added and solution became murky white. After 15 min at 0 C, the solution became a murky pink color, with a white precipitate above the solvent. Solution allowed to rise to r.t. and stirred for another 30 min. HPLC reported a new, less-polar peak ($t_R$ = 12.54 min) with UV maxima at 299 and 326 nm, consistent with Fmoc group and caged AMC, respectively. The reaction mixture was diluted with ethyl acetate, washed with HCl and $NaHCO_3$. Aqueous layer extracted with ethyl acetate and the combined organic fractions were dried over $Na_2SO_4$ and concentrated *in vacuo* to give a crude white solid, used directly in next step.

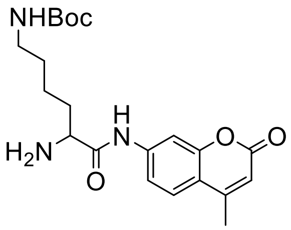

**tert-butyl (5-amino-6-((4-methyl-2-oxo-2H-chromen-7-yl)amino)-6-oxohexyl)carbamate (4):** Synthesized by an adapted literature procedure[5,6]. The starting material (6.5 mg, 0.0103 mmol) was dissolved in 2.85 mL DMF, and piperidine (15 µL) added to give 5% piperidine/DMF solution. After 5 min, TLC confirmed successful reaction via staining with ninhydrin (deep purple spot observed). Reaction diluted in ethyl acetate and water added. Extracted with ethyl acetate, shaking gently to avoid emulsions; full extraction confirmed by TLC, detecting purple stain by ninhydrin to determine product location. Organic layer dried over $Na_2SO_4$, concentrated *in vacuo* to give light-brown/yellow liquid residue, which was reconstituted in a small volume (~1-2 mL) of 50/50 MeOH/$H_2O$ for HPLC separation. Crude reside purified by FCC in 5% MeOH/DCM with 1.0% $NH_4OH$ to acquire pure product ($R_f$ = .28), as a clear solid, 2.7 mg, **65%** yield. The $^1H$ NMR spectrum was consistent with those previously reported.

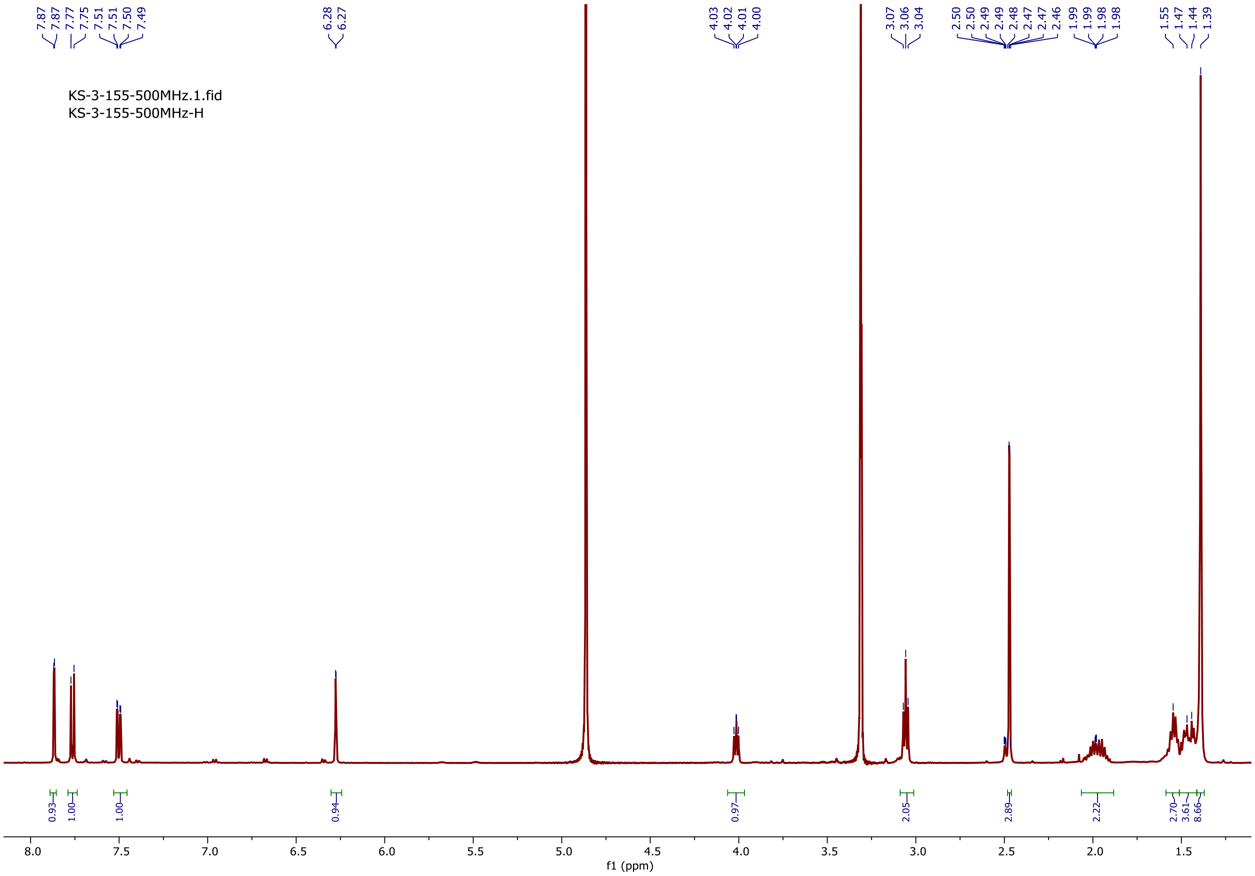

**Supporting Figure S10.** $^1H$ NMR of compound **4**.

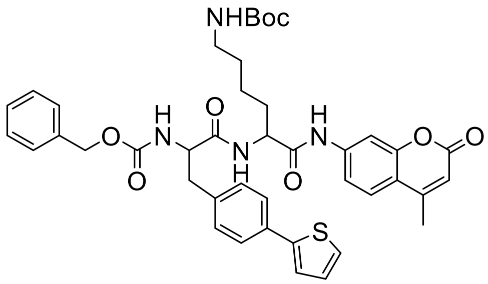

**tert-butyl (5-(2-(((benzyloxy)carbonyl)amino)-3-(4-(thiophen-2-yl)phenyl)propanamido)-6-((4-methyl-2-oxo-2H-chromen-7-yl)amino)-6-oxohexyl)carbamate (5):** Compound **2** (2.3 mg, 5.94 umol, 1.1 eq.) dissolved in DMF (2 mL), HOBt (2.4 mg, 16.2 umol, 3.0 eq.) added, then HBTU (6.0 mg, 16.2 umol, 3.0 eq.) added. Stirred for 20 min, after which compound **4** (2.2 mg, 5.4 umol, 1.0 eq.) was added. After 40 min, DIPEA (2.8 uL, 16.2 umol, 3.0 eq.) was added and pH = 9. After 1 h, HLPC indicated formation of a new product ($t_R$ = 12.41 min) with desired UV trace. Reaction diluted with ethyl acetate and water. Extracted with ethyl acetate, then the combined organic layer was washed with water. Organic layer dried over $Na_2SO_4$, concentrated *in vacuo* to give white-yellow powder, used directly in next step.

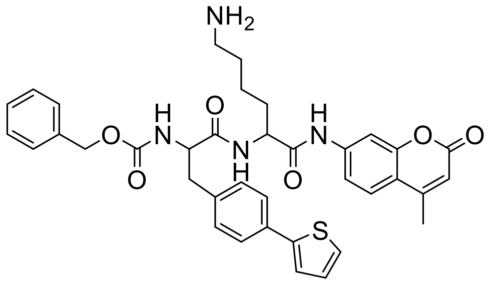

**benzyl (1-((6-amino-1-((4-methyl-2-oxo-2H-chromen-7-yl)amino)-1-oxohexan-2-yl)amino)-1-oxo-3-(4-(thiophen-2-yl)phenyl)propan-2-yl)carbamate (CTLAP):** Compound **5** (4 mg, 5.4 umol) was dissolved in DCM (2.4 mL) and trifluoroacetic acid (600 uL) was added to give a 20% TFA/DCM solution. Triisopropylsilane (1 uL, 5.4 umol, 1.0 eq.) added. After 30 min, reaction was translucent grey in color. HPLC indicated new peak ($t_R$ = 11.56 min), more polar than starting material ($t_R$ = 12.41 min). Solvent removed *in vacuo*; the resulting yellow-brown residue suspended in MeOH/$H_2O$ and centrifuged to give faint yellow solution and yellow/brown pellet. The supernatant was purified by HPLC and lyophilized to give a white powder product, 4.2 mg (**quant.**). MS: 667.2507 m/Z observed; [M+H] = 667.2582 calculated.

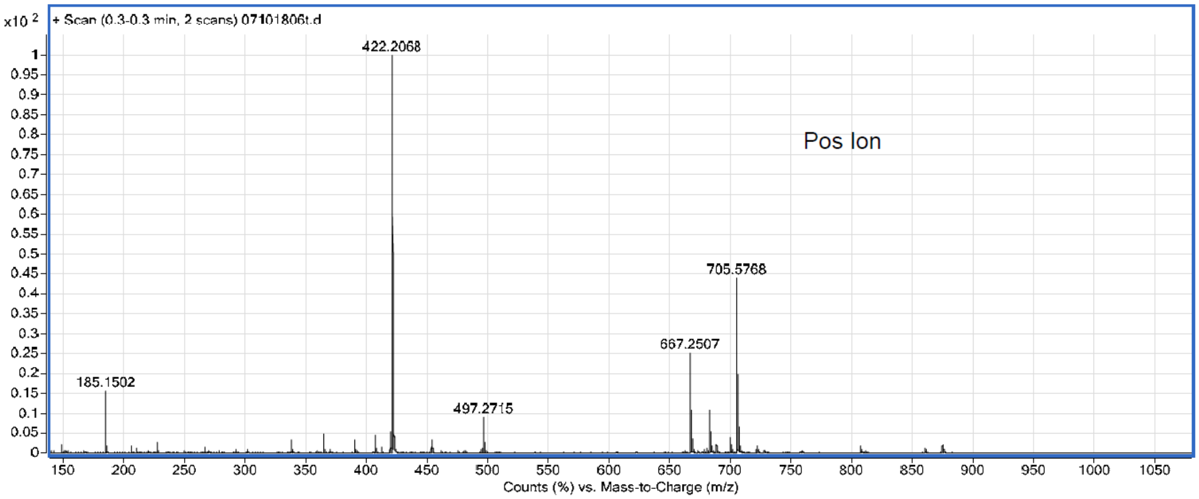

**Supporting Figure S11.** ESI-MS (negative mode) of **CTLAP**.
